# Estimating population range distributions from animal tracking data

**DOI:** 10.1101/2025.09.02.673746

**Authors:** Gayatri Anand, Christen H. Fleming, Ananke G. Krishnan, Clayton T. Lamb, E. Patrícia Medici, Laura R. Prugh, Justin M. Calabrese, William F. Fagan

## Abstract

Quantifying the space requirements of a population is a fundamental problem in spatial ecology, particularly as it relates to the identification of important utilization areas and the designation of protected areas for conservation and wildlife management. Traditionally, population space use estimation techniques scale up from the individual to the population level by aggregating individual animal tracks and then using a single pooled distribution estimator like minimum convex polygons (MCP) or kernel density estimation (KDE). These techniques fail to account for the high levels of temporal autocorrelation in modern tracking datasets, and estimates are often sensitive to the number of individuals sampled. We introduce a new population kernel density estimator (PKDE) that accounts for temporal autocorrelation in tracking data, propagates uncertainty from the individual to the population level, accounts for inter-individual variation when scaling up to the population level, and is not highly sensitive to the number of individuals tracked. Through a combination of simulated data and empirical GPS tracking datasets from three species: (a) grizzly bear (*Ursus arctos horribilis*); (b) lowland tapir (*Tapirus terrestris*); and (c) bobcat (*Lynx rufus*), we demonstrate that PKDE produces minimally biased estimates of population-level space use compared to conventional methods like MCP and KDE. The use of conventional estimators can lead to substantial underestimation of population space usage, making them unsuitable for area-based conservation planning. The statistically efficient PKDE estimator provides relatively unbiased estimates of space use with fewer individuals sampled. This method has been made available as a function in the *ctmm* R package.

## Introduction

Estimating the amount of space occupied by a population of animals, often called population range estimation, is an important goal at the interface of spatial ecology, behavior, and wildlife management. Related topics, including a species’ minimum area requirement (Shaffer 1981) and critical patch size (Skellam 1951) underpin the establishment of marine protected areas (Baskett et al. 2007; Lockwood et al. 2002; Medina-Vogel et al. 2008) and other areas designated for species conservation (Brown and Crone 2016; Lindsey et al. 2011; Pe’er et al. 2014). Increasingly, individual-level animal tracking data are playing a key role in population-level decision-making in the delineation of important conservation areas (e.g., Marine Protected Areas: Hays et al. 2019; Kramer and Chapman 1999; Important Bird Areas: Arcos et al., 2012; López-López et al., 2007; Ramirez et al., 2017; Soanes et al., 2016; and other protected areas: Chundawat et al. 2016); the identification of migratory corridors (Kauffman et al. 2021); and the monitoring of wildlife crossings (Bastille-Rousseau et al. 2018; Hardy et al. 2003; Neumann et al. 2012).

When making the leap from individual-level tracking data to population-level distributions for direct, real-world planning purposes, it is essential that the methods employed are robust, representative, and reliable. Traditionally, population range estimation in the context of conservation applications has involved combining all individuals’ movement tracks into a single dataset and then using a single pooled estimator like minimum convex polygons (MCP) or kernel density estimation (KDE) without distinguishing between individuals or accounting for temporal autocorrelation (Chandler et al. 2019; Donovan et al. 2017; Gutowsky et al. 2015; Hyrenbach et al. 2006; Obbard et al. 2010; Schaefer et al. 2008; Schaefer and Mahoney 2007; Scott et al. 2012; Starking-Szymanski et al. 2018; Watson et al. 2003). While simple to implement, this approach is problematic because it yields population-level results that ignore non-independence within individuals and are biased toward individuals with more data.

Although MCP and KDE long served as the workhorses of individual-level home range estimation, they have been shown to be ill-suited for modern tracking data because they assume independence among the data points (Noonan et al., 2019). The accurate estimation of individual home ranges from modern GPS tracking data requires statistical methods that can account for the high levels of temporal autocorrelation that these data routinely exhibit (Calabrese et al., 2016; Silva et al., 2022). When tracking data violate the assumption of being independent and identically distributed (IID)—which is almost always the case for modern GPS data—MCP and KDE estimates of home range will be negatively biased, sometimes quite badly (Börger et al. 2006; Fleming and Calabrese 2017; Noonan et al. 2019; Walter et al. 2015). This underestimation of individual-level home ranges is expected to propagate to the population level, which suggests that MCP- and KDE-based population range estimates will tend to be negatively biased. An additional challenge is that these estimation techniques are not conventionally accompanied by a corresponding uncertainty estimate, and are therefore unequipped to identify instances where range estimates have very wide confidence intervals, where further data collection might be required to obtain more precise estimates (Fleming et al. 2015).

Conventional population range estimation techniques also suffer from a second source of bias that arises from ignoring inter-individual variation when scaling up to the population level.

Specifically, conventional methods assume that the sampled individuals’ movements are representative of the entire population, and this also leads to the underestimation of the population range and its uncertainty (Gutowsky et al. 2015). A third problem is that estimates of population range size generated using MCP or KDE methods are sensitive to sample size, with the population range size tending to increase as the number of individuals sampled increases, until the estimate begins to saturate as the sample more adequately captures the population variation (Gutowsky et al. 2015; Sequeira et al. 2019; Soanes et al. 2013). Despite this negative bias associated with small population sample sizes, few studies using these estimators have used saturation plots of population range area vs. sample size to check if they have sufficiently sampled the population (but see Anderson et al., 2019; Thums et al., 2018).

Extending the widely agreed-upon quantitative interpretation of a home range (Burt 1943), we formally define a population range to be a percentage coverage area of the probability distribution of all possible locations used by the population. Building upon statistical methods previously developed for the estimation of individual-level home ranges with autocorrelated data, here we propose a new Population Kernel Density Estimator (PKDE) that calculates more accurate estimates of the population range from a small number of tracked individuals. Using a hierarchical model, PKDE estimates the population distribution mean and variance parameters from individual home range distributions. We show that PKDE outperforms conventional methods for estimating the population range by (1) accounting for temporal autocorrelation in the data, (2) propagating uncertainty to the population level to provide reliable confidence intervals, (3) accounting for variation in individual-level space use through a hierarchical model, and (4) remaining relatively insensitive to the number of individuals sampled. We compare the performance of the new method to that of conventional methods on both simulated data and empirical case studies of populations of grizzly bear (*Ursus arctos horribilis*), lowland tapir (*Tapirus terrestris*), and bobcat (*Lynx rufus*).

## Methods

### Statistical development of the PKDE approach

Given animal tracking data **r***_k_*(*t_i_* ) = (*x_k_*(*t_i_*), *y_k_*(*t_i_*)) in two dimensions on *m* independent individuals indexed by *k* and sampled at *n_k_* times *t* indexed by *i*, assuming movement is represented by a stationary process where the mean and autocorrelation structure lack temporal variation, we consider a population-weighted kernel density estimate of the form

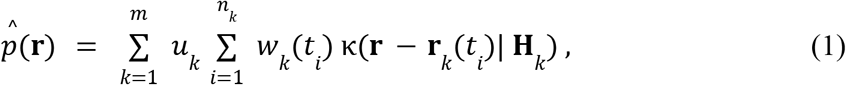

under the constraints

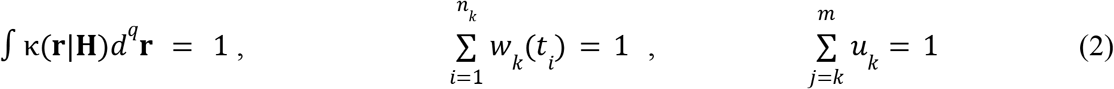

where *q* is the number of spatial dimensions of **r** (here, two). We take the kernel function κ to be Gaussian with bandwidth **H***_k_* for individual *k*. The temporal weights assigned to individual *k*, represented by *w_k_*(*t_i_*), prevent bias due to oversampled times (Fleming et al. 2018), and the weights assigned to each individual, represented by *u_k_*, prevent bias due to unequal sampling effort between individuals. The constraints in Eq. 2 ensure that the density is positive and integrates to 1, and that the kernel weights remain positive and sum to 1.

In kernel density estimation, the bandwidth is optimized by minimizing the mean integrated square error under various approximations (MISE; Fleming et al., 2018; Silverman, 1986; Turlach, 1993). Here, we perform this optimization by approximating the true population density using a Gaussian reference function estimated from a hierarchical model of individual-level Gaussian stochastic processes described in Appendix S1: Section S1. We further derive the expression for non-stationary PKDE in Appendix S1: Section S3, and a finite population correction in Appendix S1: Section S4.

### Four alternative methods for estimating population ranges

Our analyses of population range estimation compare four methods. Firstly, we used a minimum convex polygon method (MCP) implemented using the R package *adehabitatHR* (Calenge, 2011). In this approach, MCP was used to estimate the population range using tracking data that were stripped of individual identity and pooled (Chandler et al. 2019; Donovan et al. 2017; Obbard et al. 2010; Rosatte 2017; Rozylowicz and Popescu 2013; Schaefer et al. 2008; Schaefer and Mahoney 2007; Scott et al. 2012; Starking-Szymanski et al. 2018; Watson et al. 2003). This technique does not account for temporal autocorrelation in data, ignores inter-individual variation when scaling up to the population level, and is sensitive to the number of individuals sampled.

Secondly, we used a debiased Gaussian reference function (GRF) based kernel density estimation (KDE) method that assumes IID data (Fleming and Calabrese 2017), implemented using the *ctmm* **R** package on the pooled tracking data. We selected this method over more traditional KDE bandwidth selection techniques like *h*_ref_ (Silverman 1986) to account for confounding factors (e.g., differences between smoothers) and eliminate other differences in implementation. Our chosen approach allows us to isolate the effects of not modeling autocorrelation when comparing different techniques. We also used more traditional *h_ref_* bandwidth selection (implemented in *adehabitatHR*) to show that the qualitative results do not change whether we use GRF KDE in *ctmm* or *h*_ref_ in *adehabitatHR* (Appendix S1: Section S6). Traditional KDE is widely used to estimate population range utilization distributions (UDs) using pooled tracking data (Anderson et al. 2019; Bauduin et al. 2016; Chandler et al. 2019; Gutowsky et al. 2015; Hyrenbach et al. 2006; McFarlane Tranquilla et al. 2013; Morera-Pujol et al. 2023; Ra et al. 2008; Thiebot et al. 2011; Watson et al. 2003). This method of population range estimation, like MCP, also ignores temporal autocorrelation in the data and does not extrapolate out to the rest of the population.

The third technique we compared is an autocorrelated kernel density estimation (AKDE; Fleming et al., 2015) method involving the `mean` function of the **R** package *ctmm*, hereafter called the ‘Mean-AKDE’ method. In this method, we fit a series of stochastic process models for animal movement to individual movement tracks using the `ctmm.select` function in *ctmm* (Calabrese et al. 2016; Fleming et al. 2015), followed by the `akde` function with the AIC-best movement model (Fleming et al. 2015) to estimate individual-level UDs and home range contours. Lastly, we feed the individual AKDE estimates into the `mean` function of *ctmm*, which estimates the population range by generating a weighted average of the individual AKDEs. This method is an improvement over MCP and traditional KDE as it addresses the strong biases associated with the temporal autocorrelation present in modern GPS tracking data. Additionally, the *ctmm* framework allows for the incorporation of location error and non-standardized sampling schedules, both of which can introduce biases if not accounted for (Fleming et al. 2018, 2020). Even though the Mean-AKDE method represents an advance over traditional methods, it does not extrapolate out to the rest of the population, meaning that accounting for autocorrelation alone is insufficient to produce unbiased estimates.

To address the remaining problem, we develop a fourth and final technique, the PKDE method, which uses the `pkde` function of *ctmm* to return a population UD (Appendix S1: Fig. S1). Like the Mean-AKDE method, the new PKDE approach uses standard methods in *ctmm* to generate AKDE estimates of individual home ranges. However, unlike the Mean-AKDE method, the new PKDE method optimizes a population-level bandwidth to minimize the error in the population-level density estimate (Appendix S1: Eq. S1). Along with accounting for temporal autocorrelation, the PKDE technique uses a hierarchical model to account for inter-individual variation in space use. It is thus relatively insensitive to the number of individuals sampled.

In Appendix S1: Section S6, we also consider other less commonly used methods for population range estimation, such as MCP and KDE with the (unpooled) individual-level movement data, as well as “occurrence distribution” estimators (like Brownian Bridge) that are often mistakenly used as range estimators (Alston et al. 2026).

### Comparisons among methods using simulations

Researchers frequently investigate how many individuals need to be tracked to make inferences about population-level space use. Typically, studies show that the estimate of population-level space use increases with the number of individuals sampled, and asymptotes as the sample more adequately captures the population variation (Gutowsky et al. 2015; Sequeira et al. 2019; Soanes et al. 2013). We adopt the same approach here in our comparative analyses, focusing on the estimated minimum number of individuals that need to be sampled to make reliable population-level inferences.

To show the effect of sample size on the predicted population range, we first performed simulations where the ground truth (i.e., the true population range) is known. Movement data were simulated based on the continuous-time stochastic Ornstein-Uhlenbeck-F (OUF) process (Fleming et al. 2014), which models correlated positions and velocities as well as restricted space use, and is representative of modern high-quality GPS tracking data (Noonan et al. 2019). We repeatedly simulated movement tracks for a hypothetical population of 250 individuals, where the individual home range sizes were exactly *R=*1/250th the size of the population range. To do this, we assume that the home range centers are normally distributed with the mean zero and variance 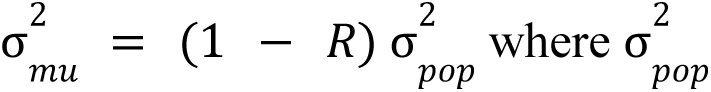 is the variance of the true population distribution and the complementary term, 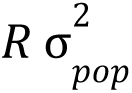, represents the variance in the individual home ranges. For each individual, movement tracks were simulated with an hourly sampling schedule for a 10-day duration. The home range crossing time and the velocity autocorrelation timescale were both set to 12 hours. Keeping the population range size and individual home range sizes fixed, we simulated movement tracks for sample sizes ranging from 10 to 250 individuals with 100 simulation runs per sample size. We then compared the performance of the four estimators on these data to the known true value. We also compared the computational complexity of different population range estimators on the simulated data for different numbers of individuals sampled (Appendix S1: Section S7). Additionally, we performed further simulations where the individual home ranges in a population were non-overlapping instead of normally distributed (Appendix S1: Section S8).

### Comparisons among methods using empirical tracking datasets

We next used GPS data from three species to demonstrate the efficacy of the new PKDE method (Appendix S1: Table S1): grizzly bears (*Ursus arctos horribilis)*, lowland tapirs (*Tapirus terrestris)*, and bobcats (*Lynx rufus)*. These species were chosen due to the duration and quality of tracking data available, and because of their tendency to remain relatively range resident which is a precondition for population range estimation. Additionally, these species vary greatly in the average degree of overlap between individuals’ ranges (Appendix S1: Fig. S6), which controls the level of bias in population range estimates (Soanes et al. 2013). As tracking data for individuals was available for different lengths of time, we partitioned the data into 1-year periods to standardize the analysis. To test the effect of sample size on the population range estimates, we randomly resampled individuals. We also tested for any effects of sex-specific differences in the movement behaviors of sampled individuals on range estimates (Appendix S1: Section S5).

We assessed the effect of sample size on the performance of the different estimators using cross-validation, where resampled training individuals were used to estimate the population range, and calculated the percentage of 200 randomly sampled GPS fixes from a test individual (not belonging to that training sample) that fell inside the predicted range. We plotted population range size as a function of sample size and compared results across all four estimators.

## Results

### Estimator performance on simulated data

The population range predicted by PKDE is larger and the contours are smoother compared to the other estimation techniques whose contours are more tightly fit to the simulated individuals’ autocorrelated tracks (Fig. 1).

**Fig 1:**
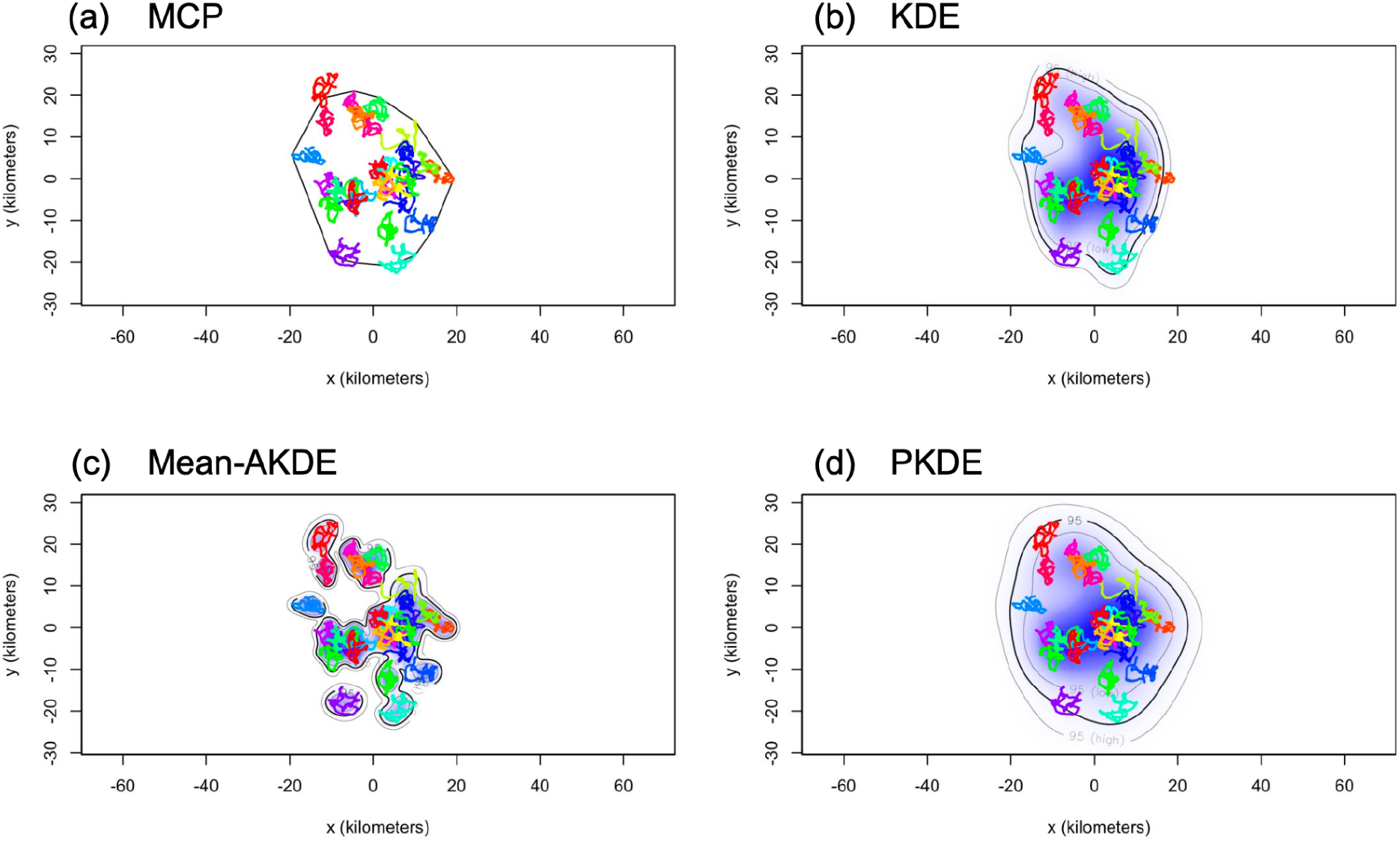
Population range of 30 simulated individuals (tracks indicated in different colors) estimated using four different techniques: (a) MCP, (b) KDE, (c) Mean-AKDE, and (d) PKDE. The thick black line represents the 95% population range estimates, the lighter contours in (b), (c), and (d) represent the 95% confidence intervals of the estimate, and the blue shading in (b), (c), and (d) represents the density estimate. The PKDE range in panel (d) is the largest of the four estimates compared here, with significant uncertainty associated with the population range estimate unlike the KDE and Mean-AKDE estimates in (b) and (c) which have smaller uncertainty. In panel (a), MCP has no associated uncertainty estimate.

On comparing the ranges predicted using the simulated data where the true population area is known, the range sizes produced by all four estimators increased with sample size, but PKDE asymptoted to the true value at smaller population sample sizes (Fig. 2). MCP, KDE, and Mean-AKDE underestimated the population range at smaller sample sizes to a much larger degree than PKDE.

**Fig 2:**
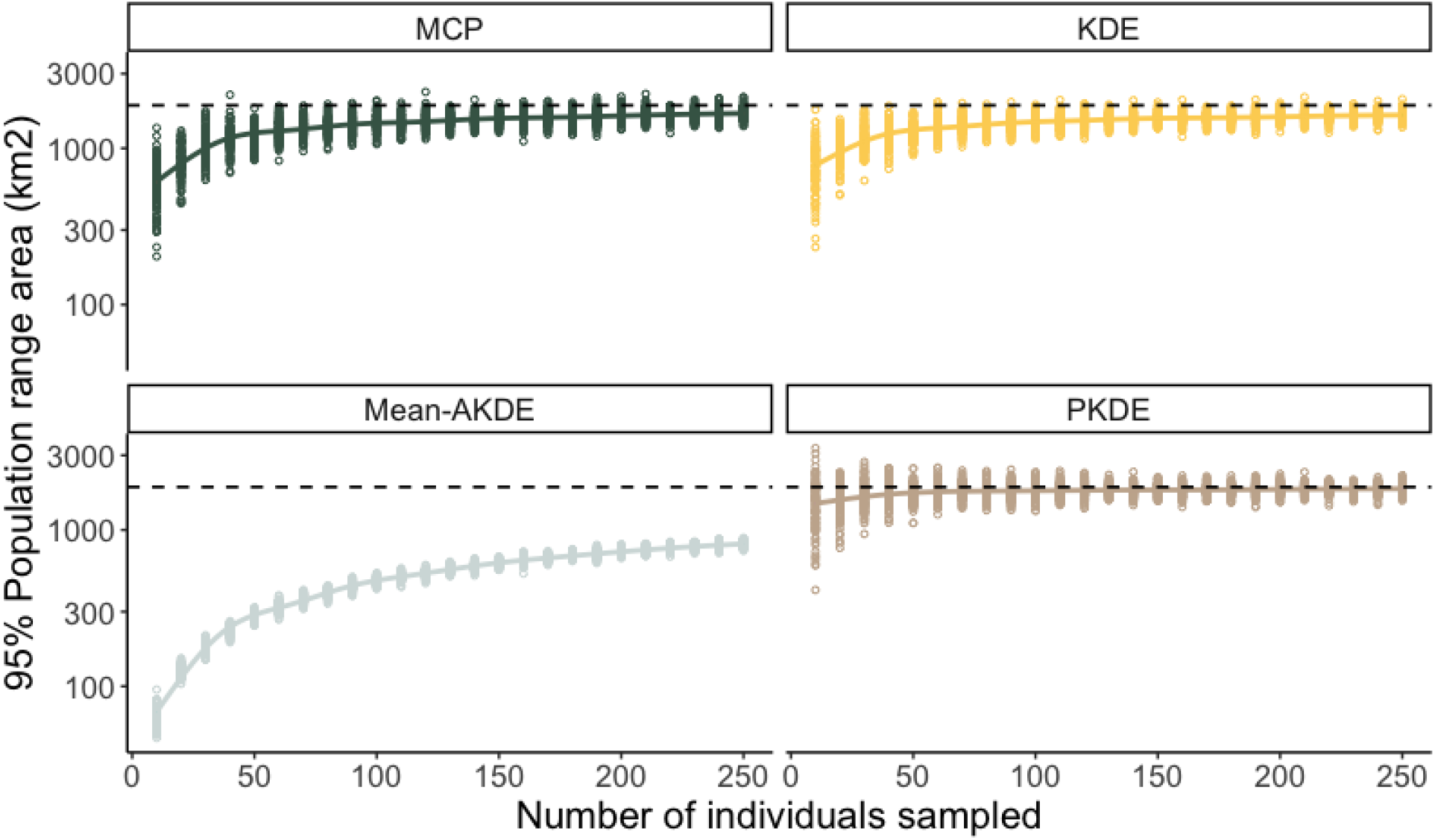
Validation of population space use estimators using simulated movement tracks where the true population range area is known. Effect of sample size on the range (km^2^) of a simulated population of 250 individuals is predicted using MCP, KDE, Mean-AKDE, and PKDE. The y-axis is log-scaled. Dashed line indicates the true population range area. Each colored open circle represents one simulation run, with 100 simulation runs per sample size group.

### Estimator performance on empirical data

The predicted range for all three species as estimated by MCP, KDE, and Mean-AKDE all increase in size with increasing sample size, as the curve asymptotically saturates (Fig. 3 a-c). The curves depicting PKDE show a similar trend, however, they asymptote at smaller population sample sizes compared to the other estimators. The PKDE estimates have a small sample size (negative) bias, however this bias is much smaller when compared to the other estimators. Additionally, at lower sample sizes, the PKDE estimates have a large variance that decreases with increasing sample size to produce more precise estimates of space use. The population range estimate derived using MCP, KDE, and Mean-AKDE for grizzly bears corresponding to the largest sample size for a given year (N=16) is comparable to the PKDE estimate at that same sample size (Fig. 3 a). However, this is not the case for the other two species: the MCP, KDE, and Mean-AKDE range estimates for tapirs and bobcats at the largest sample sizes are much smaller than the corresponding PKDE estimates (Fig. 3 b-c).

**Fig 3:**
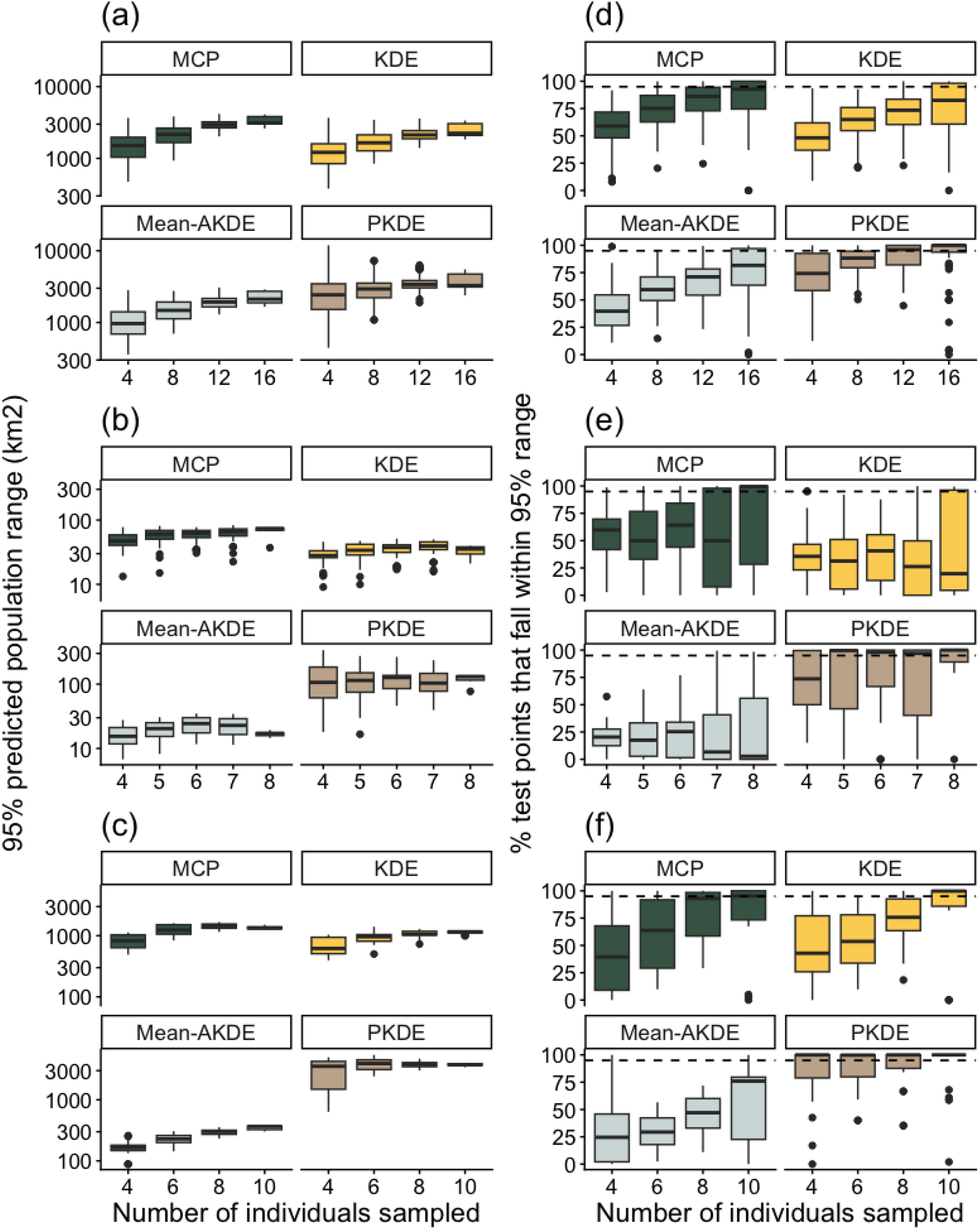
Log-linear plots of the predicted 95% population range size in km^2^ (a - c) and cross-validation analyses of 95% population UD (d - f) as a function of sample size for three empirical tracking datasets of grizzly bears (a, d), tapirs (b, e), and bobcats (c, f) subset by year, as predicted by MCP, KDE, Mean-AKDE, and PKDE. The PKDE estimates converge much faster than the other methods. In plots (d - f), the percentage of GPS fixes from unsampled individuals that fall within the 95% population range (indicated by the dashed line) predicted from a sample of individuals is plotted against the number of individuals sampled.

To evaluate the performance of the PKDE estimator, we cross-validated whether the population range (as predicted from resampled individuals’ movement data) encompassed tracks from individuals that were held out in the prediction process (Fig. 3 d-f). If the estimates reflect the population range accurately, the percentage of hold-out points that fall within the predicted population range should approximately be equal to the quantile for that UD (i.e., the 95% range estimate should ideally include 95% of the hold-out cross-validation points). The 95% UD validation curves demonstrate PKDE’s superior performance over other estimation techniques considered here. As the number of individuals sampled increases, the percentages of left-out points inside the ranges predicted by MCP, KDE, and Mean-AKDE also increase until, eventually, the curves begin to asymptote and sampling more individuals does not substantially change this value. Unlike PKDE, these methods have a consistent negative bias across lower sample sizes. The 50% and 25% UD validation curves (Appendix S1: Fig. S4 g-l) show that PKDE has minimal negative bias compared to more traditional estimators we considered, and also show that PKDE is not overestimating the population range either.

## Discussion

We have introduced a statistically efficient and asymptotically consistent estimator of the population range and demonstrated that it outperforms conventional estimators, which produce negatively biased estimates. The older MCP and KDE methods, as well as the more contemporary Mean-AKDE method, all underestimated population range considerably. Indeed, even the conventional estimates derived using the largest sample sizes possible were still smaller (often substantially smaller, as in the case of tapirs and bobcats), than the corresponding PKDE estimate. We also demonstrated that the predicted PKDE population ranges show better performance during cross-validation when compared to the other techniques. Thus, using conventional estimators for area-based conservation efforts can lead to substantial underestimation of the space requirements of a population. Additionally, unlike conventional estimators, PKDE provides more appropriate confidence intervals on its range estimates, which are useful for making informed wildlife management and conservation decisions.

The degree of space sharing in a population is expected to control the minimum number of individuals that need to be sampled in order to obtain an accurate estimate of space use using a biased estimator (Soanes et al. 2013). Theoretically, for a biased estimator, greater overlap between individual home ranges is expected to produce population range estimates that are relatively unbiased even at smaller sample sizes, whereas populations with little to no overlap will require that the entire population is sampled to obtain an accurate estimate of population space use (assuming that the data lack temporal autocorrelation, which can also be a source of negative bias). A statistically efficient estimator like PKDE is expected to produce relatively unbiased estimates, regardless of the degree of space sharing and sample size; however, the level of uncertainty associated with the estimate will be larger at lower sample sizes in scenarios with little overlap among individual home ranges. Of the species used in our empirical analysis, grizzly bears have been known to be relatively tolerant of home range overlaps (Craighead 1976; Mace and Waller 1997; Mcloughlin et al. 2000), lowland tapirs are primarily solitary (Brooks et al. 1997) but show some home range overlap (Medici 2023), and bobcats are highly territorial with females especially tending to have exclusive home ranges (Bailey 1974). We therefore expected that in the absence of temporal autocorrelation, the conventional estimators would perform best with the grizzly bear data, followed by the tapirs, and finally the bobcats. The results indicate that conventional population range estimators produce range sizes for the grizzly bear dataset that are comparable to the PKDE estimate at N=16. However, the conventional estimators considerably underestimate space use in the case of the tapir and bobcat datasets, even at the largest sample sizes possible in a given year (N=8 for tapirs, and N=10 for bobcats). This indicates that if we used the conventional estimators, further sampling of those populations would be necessary to obtain reliable estimates of the population range size. In contrast, the curves for the PKDE estimator remain relatively flat at larger sample sizes (Fig. 3 a-c).

For more than 50 years, researchers have explored whether having a small number of large protected areas or a large number of smaller protected areas is better for conservation outcomes (i.e., the single large or several small “SLOSS” debate: Diamond 1975; Wilson and Willis 1975). Different populations have different area requirements for survival (Diamond 1975), and as a consequence of the increasing availability of tracking data, home ranges have become an increasingly important component of these discussions (Allen and Singh 2016; Di Franco et al. 2018; Goldingay 2015; Laver and Kelly 2008; Nordberg et al. 2021; Zeale et al. 2012). In the cases of the tapirs and bobcats, which exhibit low levels of spatial overlap across individual home ranges, the answer to the SLOSS debate will incorrectly be determined to be several small when using poor-performing population range estimation techniques (Appendix S1: Fig. S5).

Estimators like KDE and Mean-AKDE tightly fit the population range to the sample tracking datasets and do not capture the rest of the population, producing multimodal population range distributions that favor the several small reserve design. However, for all three species, the population distributions estimated by MCP and PKDE involve undivided large ranges. This suggests that the single large reserve will produce better conservation outcomes for these populations. Although the MCP and PKDE methods yield the same result on this point, the MCP technique is not preferred because it underestimates space use in cases where the sample size is small (Fig. 3) or when there is little overlap between individual home ranges.

To make population-level inferences using conventional estimators, studies have used saturation curves to determine the minimum number of individuals that need to be sampled for reliable estimates. However, such saturation curves exhibit correlated errors (because the same total dataset is used for each subsample along the x-axis), and that correlation approaches 100% as the sampling effort reaches a maximum, making such saturation curves difficult to reliably interpret. The PKDE method of population range estimation produces accurate estimates of population space use using relatively fewer individuals in comparison to the other estimators . Obtaining a given level of accuracy with a lower sample size is beneficial because it reduces associated costs (in terms of money, effort, and threats to animal health) when determining the number of individuals that need to be tagged (Sequeira et al. 2019). Moreover, PKDE is highly adaptable and can make the most of movement data that has already been collected for other studies, potentially obviating the need for further data collection in some cases. Finally, PKDE is highly flexible and robust to different sampling methods between movement datasets, which we hope will facilitate collaborative and synthetic studies.

## Author Contribution Statement

G.A., C.H.F., J.M.C., and W.F.F. conceptualized the study. G.A. and A.G.K. performed the statistical analyses and visualizations under guidance from C.H.F. and W.F.F., with J.M.C. providing further suggestions regarding the statistical analyses. C.T.L., E.P.M., and L.R.P. provided data and biological insights. C.H.F. developed new computer code that can be accessed through the open-source *ctmm* **R** package. G.A. and W.F.F. wrote the initial draft, with C.H.F. and J.M.C. providing inputs. All authors discussed the results, contributed critically to early drafts, and gave final approval for the publication of the paper.

## Notes

### Competing Interest Statement

The authors have declared no competing interest.

### Summary of Updates

Additional analyses have been added to the Appendix, and the main manuscript has been updated to reflect this. Minor changes to the main manuscript have also been made.

