## Appendix S1 for "Estimating population range distributions from animal tracking data"

### Estimating population range distributions from animal tracking data

Gayatri Anand[1*], Christen H. Fleming[1,2,3], Ananke G. Krishnan[1], Clayton T. Lamb[4,5], E. Patrícia Medici[6,7,8], Laura R. Prugh[9], Justin M. Calabrese[1,10,11], William F. Fagan[1]

### Section S1. Minimizing the Mean Integrated Square Error (MISE)

The bandwidth and weights of the population kernel density estimate are optimized by minimizing the MISE, which expands to:

$$MISE = \int d^q\mathbf{r} \, \langle \, |\hat{p}(\mathbf{r}) - p(\mathbf{r})|^2 \, \rangle \, , \qquad \text{(Eq. S1)}$$

$$= \int d^q\mathbf{r} \, \langle \hat{p}(\mathbf{r}) \, \hat{p}(\mathbf{r}) \rangle - 2\int d^q\mathbf{r} \, \langle p(\mathbf{r}) \, \hat{p}(\mathbf{r}) \rangle + \int d^q\mathbf{r} \, \langle p(\mathbf{r}) p(\mathbf{r}) \rangle \, , \qquad \text{(Eq. S2)}$$

requiring three terms to calculate. We approximate true density, $p(\mathbf{r})$, with a Gaussian reference function

$$p(\mathbf{r}) \approx N(\mathbf{r}|\mathbf{M}, \boldsymbol{\Sigma}), \qquad \text{(Eq. S3)}$$

which we estimate from the individual Gaussian reference functions via the hierarchical model implemented in the `mean` function of *ctmm*. The population mean location, $\mathbf{M}$, is calculated from a normal meta-analysis on the individual mean locations, $\boldsymbol{\mu}_k$, using a normal sampling distribution for each mean location estimate's uncertainty and a normal population distribution for all of the individual mean locations. The population auto-covariance parameters with the population location covariance, $\mathbf{\Sigma}$, are calculated from a log-normal meta-analysis (Section S2) on the individual auto-covariance parameters with the individual location covariances, $\boldsymbol{\sigma}_k$, using a normal sampling distribution for the matrix-log transformed auto-covariance parameters' uncertainty and a normal population distribution for all of the individual log-covariance parameters. Frequentist hierarchical models are analogous to Bayesian hierarchical models, but without subjective priors. I.e., here there are only two levels to the hierarchy—the individual and the population levels. Furthermore, the use of a Gaussian reference function to approximate the true population density is known to be asymptotically optimal for kernel density estimation (Turlach, 1993). For smaller amounts of data, the Gaussian reference function KDE will be positively biased if the true distribution is not approximately unimodal. In the context of movement data, starkly multimodal data often correspond to different behaviors or populations that need to be segmented or clustered, regardless, and the resulting constituent behaviors/populations are often approximately unimodal.

### Section S2. Population location covariance calculation

We calculate the population autocovariance parameters and the population location covariance, $\mathbf{\Sigma}$, through a log-normal meta-analysis of the individual auto-covariance parameters with the individual location covariances, $\boldsymbol{\sigma}_k$, including their uncertainty (Berkey et al., 1998; Kalaian & Raudenbush, 1996). For the log-normal meta-analysis, we perform matrix-log transformations (Gantmakher, 2000; Hall, 2015) of the covariance matrices, and apply log-$\chi^2$ bias corrections (Bartlett & Kendall, 1946) after diagonalizing the log-transformed auto-covariance estimates, which provides exact bias corrections for IID isotropic processes.

### Section S3. Non-stationary PKDE derivation

Starting with the same form of data and expression for MISE introduced above in Section S1 (Eq. S1, S2), we derive a more general expression for PKDE for a non-stationary process where the mean and autocorrelation structure change over time. The basic mathematical structure for non-stationary autocorrelated kernel density estimation (AKDE) was worked out in (Fleming et al., 2015) (App. B), and here similarly resolves to

$$\int d^q\mathbf{r}\,\langle \hat{p}(\mathbf{r})\,\hat{p}(\mathbf{r})\rangle = \sum_k {u_k}^2 \sum_{i,j} \frac{w_k(t_i)w_k(t_i)}{\sqrt{(2\pi)^q det\boldsymbol{\Sigma}_{kk}(t_i,t_j)}} \qquad \text{(Eq. S4)}$$

$$+ \sum_{k\neq l} u_k \frac{e^{-\frac{1}{2}(\boldsymbol{\mu}_k-\boldsymbol{\mu}_l)^T \boldsymbol{\Sigma}_{kl}^{-1}(\boldsymbol{\mu}_k-\boldsymbol{\mu}_l)}}{\sqrt{(2\pi)^q det\boldsymbol{\Sigma}_{kl}}} u_l\,, \qquad \text{(Eq. S5)}$$

$$\int d^q\mathbf{r}\,\langle p(\mathbf{r})\,\hat{p}(\mathbf{r})\rangle = \sum_{k=1}^{m} u_k \frac{e^{-\frac{1}{2}(\boldsymbol{\mu}_k-\mathbf{M})^T \boldsymbol{\Sigma}_{k}^{-1}(\boldsymbol{\mu}_k-\mathbf{M})}}{\sqrt{(2\pi)^q det\boldsymbol{\Sigma}_{k}}}\,, \qquad \text{(Eq. S6)}$$

$$\int d^q\mathbf{r}\,\langle p(\mathbf{r})p(\mathbf{r})\rangle = \frac{1}{\sqrt{(2\pi)^q det\boldsymbol{\Sigma}}}\,, \qquad \text{(Eq. S7)}$$

in terms of the correlation functions

$$\boldsymbol{\Sigma}_{kk}(t,t') = 2\gamma_k(t,t') + 2\mathbf{H}_k\,, \qquad \text{(Eq. S8)}$$

$$\boldsymbol{\Sigma}_{kl} = \boldsymbol{\sigma}_k + \boldsymbol{\sigma}_l + \mathbf{H}_k + \mathbf{H}_l\,, \qquad \text{(Eq. S9)}$$

$$\boldsymbol{\Sigma}_{k} = \boldsymbol{\Sigma} + \boldsymbol{\sigma}_k + \mathbf{H}_k\,, \qquad \text{(Eq. S10)}$$

and where the stationary semi-variance function, $\gamma_k(t,t')$, is given by

$$\gamma_k(t,t') = \frac{1}{2}\left\langle (\mathbf{r}_k(t) - \mathbf{r}_k(t'))(\mathbf{r}_k(t) - \mathbf{r}_k(t'))^T \right\rangle = \boldsymbol{\sigma}_k - COV[\mathbf{x}_k(t), \mathbf{x}_k(t')]\,. \qquad \text{(Eq. S11)}$$

For computational efficiency, we first optimize the time weights, $w_k(t)$, for individual distribution estimation (Fleming et al., 2018), leaving us to potentially optimize the individual weights, $u_k$, and individual bandwidths, $\mathbf{H}_k$. Furthermore, optimization of the individual weights, $u_k$, for a given set of bandwidths, $\mathbf{H}_k$, is a well-defined quadratic programming (QP) problem. Therefore, our attention turns to the individual bandwidths, $\mathbf{H}_k$, for which there are two parsimonious choices that we have implemented and tested:

$$\mathbf{H}_k = h^2 \mathbf{\Sigma}\,, \qquad \texttt{kernel='population'} \qquad \text{(Eq. S12)}$$

$$\mathbf{H}_k = h^2 \boldsymbol{\sigma}_k\,, \qquad \texttt{kernel='individual'} \qquad \text{(Eq. S13)}$$

whereby the individual kernels either conform to the shape and scale of the population distribution or to the shape and scale of the individual distributions, and a single scalar bandwidth, $h$, is optimized. In our software implementation, `kernel='individual'` is the default option, but only because `kernel='population'` requires computationally expensive $n_k \times n_k$ determinant calculations in the MISE, given that $\gamma_k$ and $\mathbf{H}_k$ are not proportional. In our testing, the two methods produced quantitatively similar results over a wide range of parameter values, indicating a lack of sensitivity to this choice of bandwidth structure.

### Section S4. Finite population correction

If every member of a population is sampled, or $n_{sample} = n_{population}$, then the average of the individual AKDEs is our best estimate of the current population's distribution. On the other hand, if the $n_{sample}$ tracked individuals represent a small fraction of a much larger population, then PKDE is our best estimate of the population distribution. For intermediate cases, we can apply the finite population correction $(n_{sample}/n_{population})\langle AKDE(x,y)\rangle + (1 - n_{sample}/n_{population})PKDE(x,y)$, which interpolates between the mean-AKDE and PKDE density functions to obtain an estimate of the finite population's current distribution. However, in many situations, users may be interested in extrapolating to future generations of a population, which PKDE estimates without the finite population correction.

### Section S5. Effect of sex-specific differences on PKDE

In the case of species with sex-specific differences in movement behaviours and home range size, it is possible that the sex ratio of the individuals sampled will affect the population range size estimate. The sexes of the individuals sampled for each species for each year are detailed in Table S1.

**Table S1:** The total number of individuals sampled for each species for a given year, including a breakdown of how many of those sampled individuals are male or female.

| **Species** | **Population location** | **Year** | **Total number of individuals sampled** | **Number of males sampled** | **Number of females sampled** |
|---|---|---|---|---|---|
| Grizzly bear *(U.a.horribilis)* | British Columbia, Canada | 2017 | 14 | 5 | 9 |
| Grizzly bear *(U.a.horribilis)* | British Columbia, Canada | 2018 | 18 | 7 | 11 |
| Grizzly bear *(U.a.horribilis)* | British Columbia, Canada | 2019 | 17 | 5 | 12 |
| Grizzly bear *(U.a.horribilis)* | British Columbia, Canada | 2020 | 17 | 4 | 13 |
| Grizzly bear *(U.a.horribilis)* | British Columbia, Canada | 2021 | 12 | 1 | 11 |

| | | | | | |
|---|---|---|---|---|---|
| Grizzly bear *(U.a.horribilis)* | British Columbia, Canada | 2022 | 10 | 4 | 6 |
| Bobcat (*Lynx rufus*) | Washington, USA | 2019 | 5 | 2 | 3 |
| Bobcat (*Lynx rufus*) | Washington, USA | 2020 | 11 | 5 | 6 |
| Bobcat (*Lynx rufus*) | Washington, USA | 2021 | 5 | 2 | 3 |
| Lowland tapir (*Tapirus terrestris*) | Pantanal, Brazil | 2014 | 8 | 3 | 5 |
| Lowland tapir (*Tapirus terrestris*) | Pantanal, Brazil | 2015 | 7 | 4 | 3 |
| Lowland tapir (*Tapirus terrestris*) | Pantanal, Brazil | 2016 | 6 | 4 | 2 |
| Lowland tapir (*Tapirus terrestris*) | Pantanal, Brazil | 2017 | 8 | 3 | 5 |
| Lowland tapir (*Tapirus terrestris*) | Pantanal, Brazil | 2018 | 9 | 5 | 4 |

On comparing the differences in AKDE home range sizes between males and females for each of the three focal species, we find a significant difference for all three species for certain years (Fig. S8). In order to further explore the effect of sex ratio of sampled individuals on the PKDE population range estimate, we compared PKDE area estimates for sampled females only for a given year to the PKDE area estimates for all individuals sampled in a given year (Fig. S9). We could not include the PKDE area estimates for sampled males only as for grizzlies in 2021, only 1 male individual was sampled, and this is too low of a sample size for PKDE. We found no significant differences between the population range size estimated using the data from sampled females only, compared to data from all sampled individuals for all three species.

For species that have sex-specific differences in movement, we recommend analyzing the effect that the sex ratio of sampled individuals has on the estimated population range, and making sure that the sampled individuals are representative of the population.

### Section S6. Other population range estimation techniques

In addition to the four population range estimation techniques presented in the main text, including 1) MCP using pooled population data (referred to here as 'MCP Pooled'), 2) KDE using pooled population data implemented in *ctmm* (referred to here as 'KDE (*ctmm*)'), 3) Mean-AKDE, and 4) PKDE, we also include here comparisons with five other methods (Figs. S2, S3, S4, S5, S7, S11). These include the 'MCP Union' method which involves estimating MCP home ranges for each sampled individual and taking their spatial union to predict the population range. We also include two alternative KDE methods, both implemented in *adehabitatHR* using the $h_{ref}$ bandwidth selection technique. The first KDE technique, hereafter referred to as 'KDE Pooled ($h_{ref}$)', uses pooled population tracking data similar to the 'KDE (ctmm)' method, but makes use of the $h_{ref}$ bandwidth selection technique. The second KDE technique, referred to as 'KDE Union ($h_{ref}$)', involves estimating individual KDE home ranges and taking their spatial union to be the population range.

Finally, we include two 'occurrence distribution' estimators, which are often mistakenly used as range distribution estimators. Occurrence distribution techniques like Brownian Bridges (Benhamou, 2011; Benhamou & Cornélis, 2010; Horne et al., 2007) and other stochastic bridge-based techniques do not characterize an animal's or population's use of space, but instead characterize the uncertainty in their spatial position (Alston et al., 2026; Fleming et al., 2015; Signer & Fieberg, 2021), making them unsuitable for population range estimation. Despite this, we recognize that random bridges are often mistakenly used as range estimators, so two such techniques have been included in our comparative analysis: 1) Brownian Bridges (Benhamou, 2011; Horne et al., 2007; Kranstauber et al., 2014; Pozdnyakov et al., 2014) and 2) kriged occurrence distribution with the optimal movement model following model selection, where the candidate models include Brownian Motion (BM; which has continuous position but discontinuous velocity) and integrated Ornstein-Uhlenbeck motion (IOU; which has continuous position and continuous velocity, Fleming et al., 2016), hereafter referred to as 'BM/IOU Krige'. Both these methods were implemented using the *ctmm* R package. We included the BM/IOU Krige method in addition to Brownian bridges because when continuous-time stochastic movement models were fit to the empirical movement tracks for the three focal species, the BM model was misspecified in 75.9% of all cases, and the IOU model was found to be a better fit to the data. The PKDE estimation method has better performance during cross-validation when compared to these five additional population range estimation techniques (Figs. S3, S4).

Among related occurrence distribution methods, the biased random bridge (which is derived from a model of Brownian motion with linear drift) is no different in structure from ordinary Brownian Bridge (derived without drift) because the drift terms cancel out algebraically and the final result has no dependence upon the drift, as pointed out by Kranstauber et al. (2014) and as proved mathematically in Appendix S1 of Fleming et al., (2016). The only result of including drift is to shrink the diffusion-rate parameter estimate and thus shrink the resulting occurrence distribution. Furthermore, although it is not the default method employed by *adehabitatHR* (or, to the best of our knowledge, any other R package that calculates Brownian bridges), some researchers have suggested using KDE's bandwidth in the Brownian bridge (Benhamou, 2011; Benhamou & Cornélis, 2010), which produces a distribution estimate that is extremely similar to the KDE (already considered in our analysis), as optimal KDE bandwidths tend to be substantially larger than typical step lengths, unless the bandwidth itself is severely underestimated.

### Section S7. Computational Complexity of PKDE

For the population range being calculated using data from $N$ sampled individuals, each of which has $n_i$ sampled fixes, where $i = 1, \dots, N$, the computational complexity of fitting a CTMM movement model to an individual's telemetry data is $O(n)$. For AKDE home range estimation, the computational complexity of optimal bandwidth calculation can be between $O(n \log n)$ and $O(n^3)$, depending on the selected method. When estimating the PKDE population range, the computational complexity is $N$ times that of the AKDE.

In Fig. S7, we present a comparison of the computational cost in seconds (log-transformed) and mean relative error (%) of different population range estimation techniques corresponding to the simulation study introduced in the main text. We conducted 100 simulation runs for each sample size. PKDE has the highest computational cost of the 9 estimators we considered, followed closely by Brownian bridge and BM/IOU Krige. The remaining estimators (MCP Union, MCP Pooled, KDE Union ($h_{ref}$), KDE Pooled ($h_{ref}$), KDE (*ctmm*), and Mean-AKDE) had a significantly lower computational cost. However, when comparing the mean relative error of each of these estimation techniques, PKDE consistently outperforms the other estimators, especially at lower sample sizes, followed by the KDE (*ctmm*), KDE Pooled ($h_{ref}$), and MCP Pooled estimators. The worst-performing estimators are "union" estimators (KDE Union ($h_{ref}$) and MCP Union), Mean-AKDE, and the occurrence distribution estimators (Brownian Bridge and BM/IOU Krige).

Despite the fact that PKDE has a high computational cost compared to the other estimators, it offers much more reliable population range estimates, especially when the number of sampled individuals is low. This is particularly notable when considering the fact that most empirical tracking datasets include only a small fraction of the total population due to the high costs (in terms of money, effort, and threat to animals' health) associated with tagging. Although it is tempting to use estimators like KDE (*ctmm*), which has the second-lowest mean relative error after PKDE and has a relatively low computational cost, we have shown that when used with empirical tracking data, it does not cross-validate well (Fig. 3 d-f, Fig. S4), and tends to underestimate the population range (Figs. S4 and S5).

Furthermore, KDE tends to underestimate the degree of uncertainty in the population range. For example, in Fig. S5, the uncertainty estimated by the KDE method is so small that the 95% CI contours are barely visible. MCP estimates are not conventionally associated with an uncertainty estimate. PKDE propagates uncertainty to the population level to provide reliable confidence intervals, which is important because having accurate and unbiased confidence intervals of population range estimates allows for statistically sound analyses.

### Section S8. Simulations with non-overlapping home ranges

The assumption of the Gaussian distribution of home range centers made by the simulation study presented in the main text does not accurately represent territorial individuals, where the individual home ranges in a population are non-overlapping. We conducted further simulation studies to analyze and compare the performance of different population range estimators when individuals have non-overlapping home ranges. We start by fixing the home range mean locations on an 11x11 grid and assuming that all individuals have the same home range size, such that individuals have little to no overlap in their home ranges (Fig. S10). The population size was, therefore, fixed at 121 individuals. We then randomly sampled 10 to 120 individual home range centers from this 11x11 grid and simulated movement tracks for these sampled individuals centered on the selected home range centers. Movement data were simulated based on the OUF process, and the home range crossing time and the velocity autocorrelation timescale were both set to 12 hours. We conducted 100 simulation runs per sample size, and the performance of all the population range estimators was compared against the true population range area. For the PKDE estimate, we used the finite population correction detailed in S4.

On comparing the population range size predicted by different estimators to the true known population range area (Fig. S11), PKDE has some positive bias at small sample sizes; however, on increasing the sample size, the estimate converges to the true value. KDE (*ctmm*) always overestimates the population range, regardless of the number of individuals sampled. The KDE Pooled ($h_{ref}$) estimator performs well at small sample sizes, but starts overestimating population range as sample size is increased. The MCP Pooled estimator has a small negative bias at smaller sample sizes, but it quickly converges to the true population range area. Mean-AKDE and KDE Union ($h_{ref}$) significantly underestimate population range size at small sample sizes, and slowly converge to the true population range area, with estimates being comparable to the true value only at the largest sample sizes. Estimates made using the MCP Union, Brownian Bridge, and BM/IOU Krige methods increase as the sample size is increased, but always significantly underestimate the population range size.

Section S9. Supplementary figures

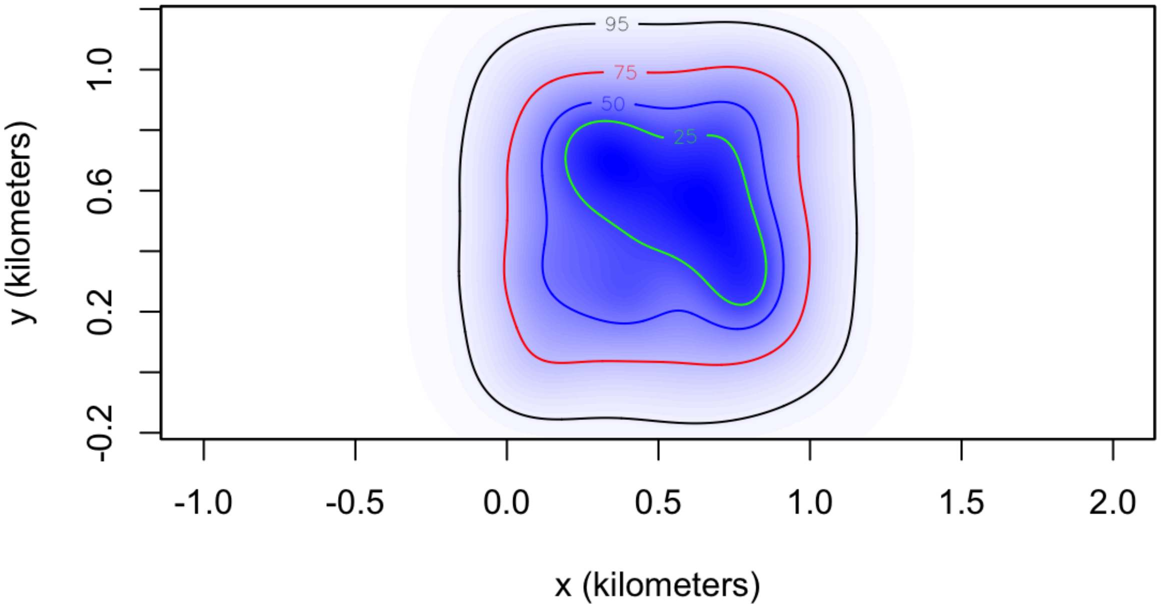


**Figure S1:** A population range distribution with the 95% (black), 75% (red), 50% (blue), and 25% (yellow) UDs boundaries indicated by the coloured solid lines. The blue shading corresponds to the density estimates.

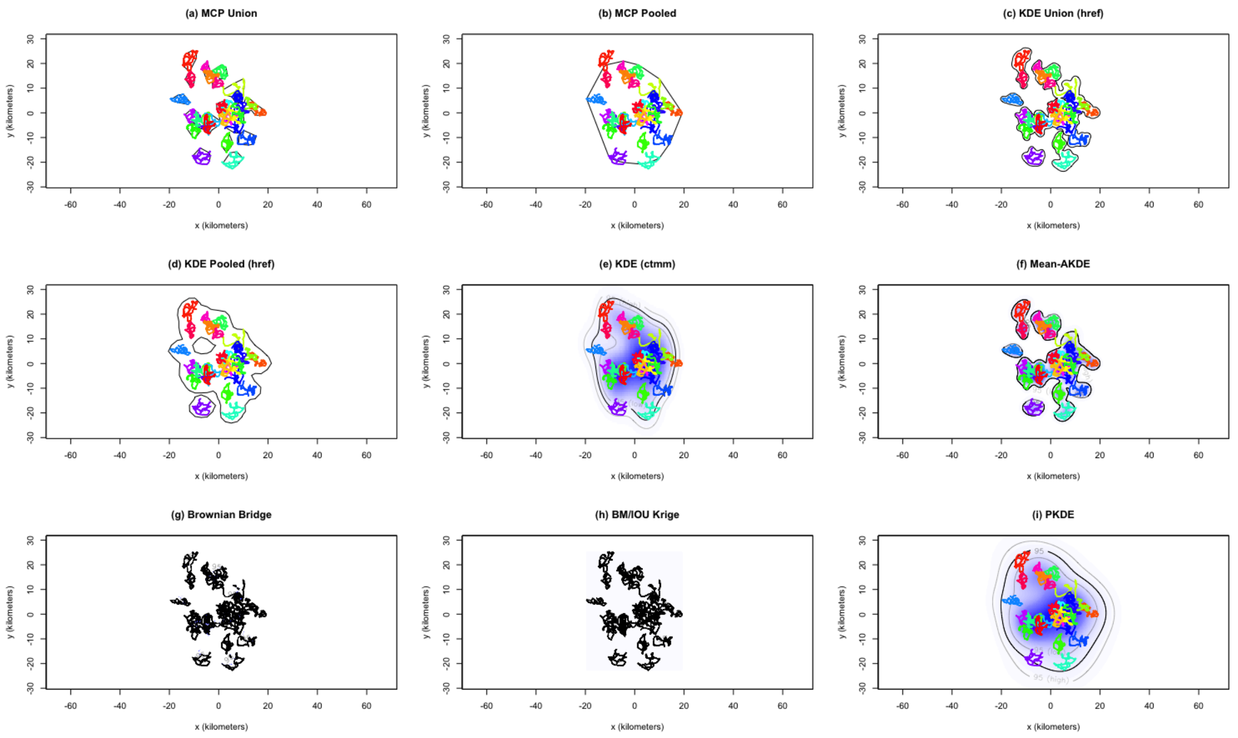


**Figure S2:** Population range of 30 simulated individuals (tracks indicated in different colors) predicting using nine different estimation techniques: (a) MCP Union, (b) MCP Pooled, (c) KDE Union ($h_{ref}$), (d) KDE Pooled ($h_{ref}$), (e) KDE (*ctmm*), (f) Mean-AKDE, (g) Brownian bridge, (h) BM/IOU Krige, and (i) PKDE. The thick black line represents the 95% population range estimates, the lighter contours in (e), (f), (g), (h), and (i) represent the confidence intervals of the estimate, and the blue shading represents the density estimates. The simulated tracks are not plotted in (g) and (f) to avoid masking the population UDs produced by Brownian Bridge and BM/IOU Krige, which fall tightly around the movement tracks.

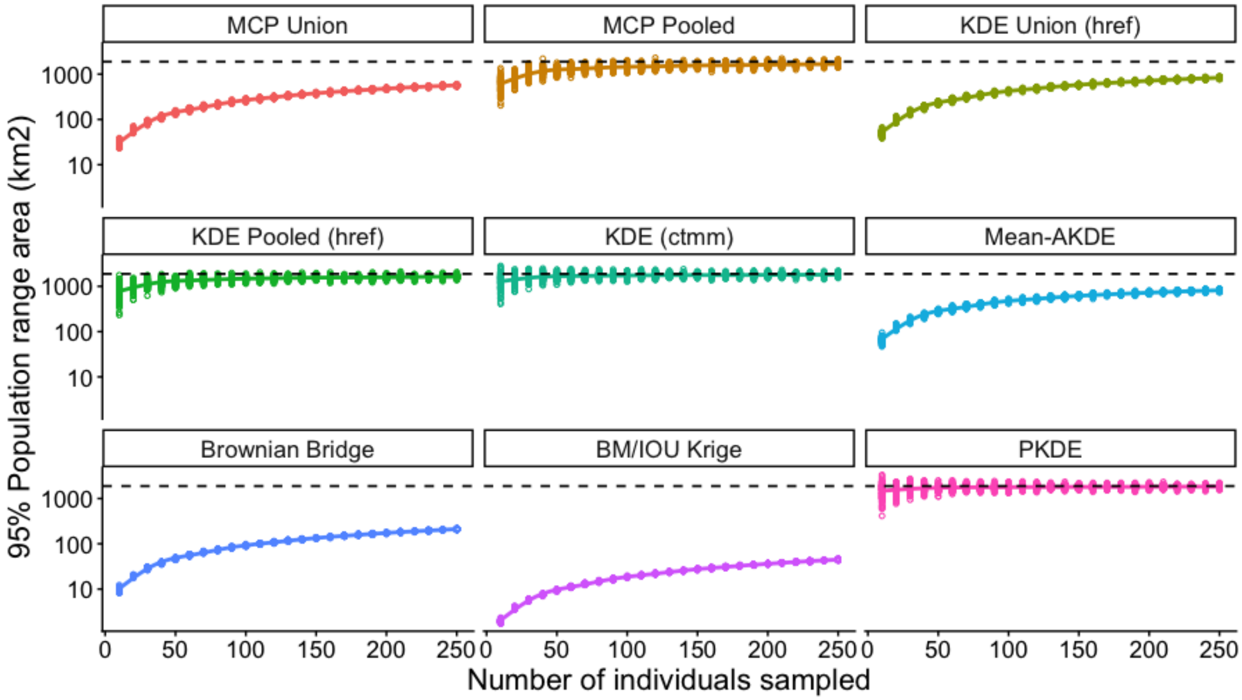


**Figure S3:** Validation of population space use estimators using simulated movement tracks where the true population range area is known. Effect of sample size on the range ($km^2$) of a simulated population of 250 individuals predicted using MCP Union, MCP Pooled, KDE Union ($h_{ref}$), KDE Pooled ($h_{ref}$), KDE (*ctmm*), Mean-AKDE, Brownian Bridge, BM/IOU Krige, and PKDE. The y-axis is log-scaled. Dashed line indicates the true population range area. Each colored open circle represents one simulation run.

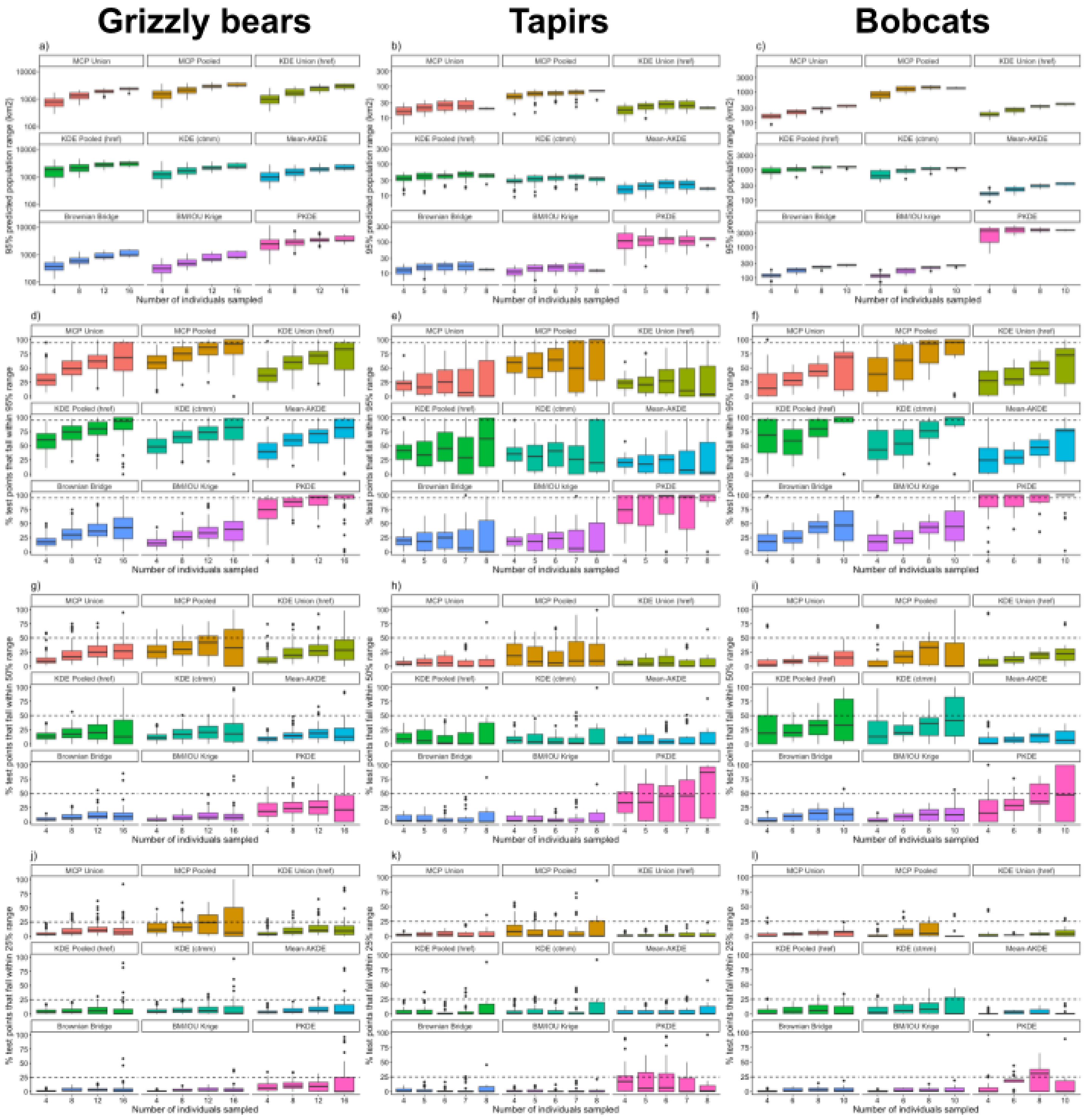


**Figure S4:** Cross-validation analyses using three empirical datasets (grizzly bears, tapirs, and bobcats) to compare the performance of nine estimators of population range size as a function of sample size. First row (a-c): Log-linear plots of the population 95% UD (km2) predicted by MCP Union, MCP Pooled, KDE Union ($h_{ref}$), KDE Pooled ($h_{ref}$), KDE (*ctmm*), Mean-AKDE, Brownian Bridge, BM/IOU Krige, and PKDE for different numbers of individuals sampled. Second row (d-f): Percentage of GPS fixes from unsampled individuals that fall within the 95% population range (indicated by the dashed line) predicted from a sample of individuals. Third row (g-i): Percentage of GPS fixes from unsampled individuals that fall within the 50% population range (indicated by the dashed line) predicted from a sample of individuals. Fourth row (j-l): Percentage of GPS fixes from unsampled individuals that fall within the 25% population range (indicated by the dashed line) predicted from a sample of individuals.

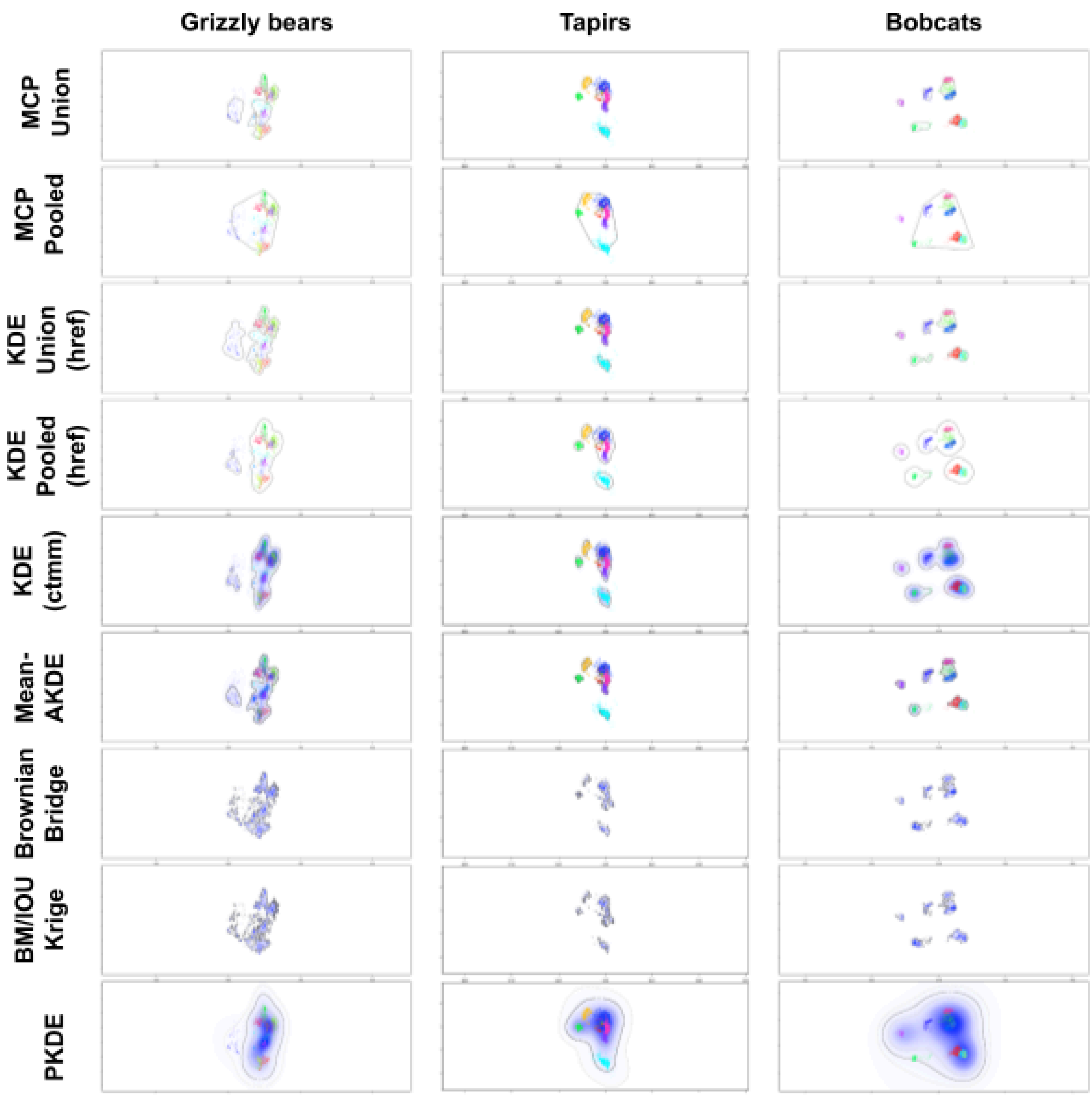


**Figure S5:** Population range of the three focal species: 1) grizzly bears (N=16, year=2018), 2) tapirs (N=8, year=2018), and 3) bobcats (N=10, year=2020), estimated using nine techniques: MCP Union, MCP Pooled, KDE Union ($h_{ref}$), KDE Pooled ($h_{ref}$), KDE (*ctmm*), Mean-AKDE, Brownian Bridge, BM/IOU Krige, and PKDE. 95% UD contours are indicated using thick black lines. Lighter lines in KDE (ctmm), Mean-AKDE, Brownian bridge, BM/IOU Krige, and PKDE plots correspond to the 95% CIs of the predicted range, and the blue shading corresponds to the density estimates. The uncertainty estimated by the KDE (ctmm), Mean-AKDE, Brownian bridge, and BM/IOU Krige methods is so small that the 95% CI contours are barely visible. The MCP Pooled range for bobcats does not include the tracks of one individual indicated in purple as that individual had only 379 GPS fixes and therefore was excluded from the 95% "core" MCP range for the population.

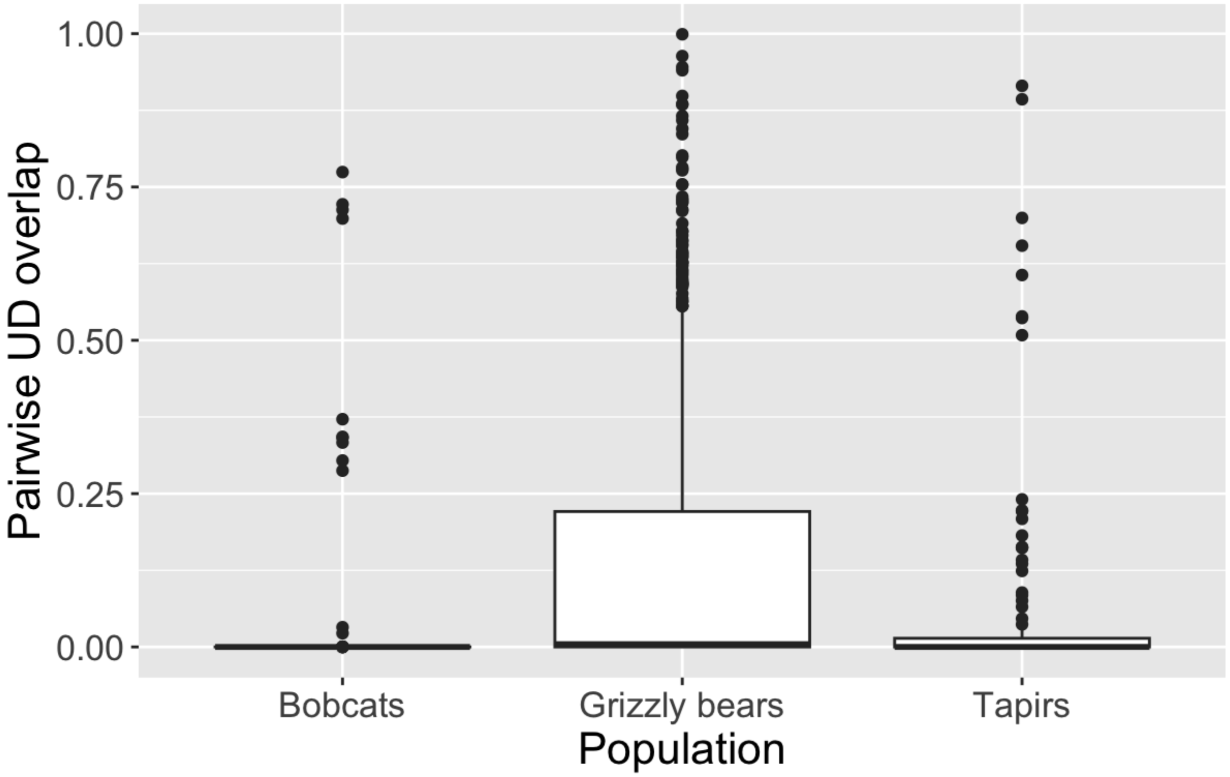


**Figure S6:** The degree of space sharing in the three focal species estimated by measuring the pairwise range overlap (Bhattacharyya coefficient; Winner et al., 2018) between individuals' home ranges estimated using autocorrelated kernel density estimation (Fleming et al., 2015). Grizzly bears are much more tolerant of home range overlaps than Lowland tapirs, which are mainly solitary, and bobcats, which are highly territorial.

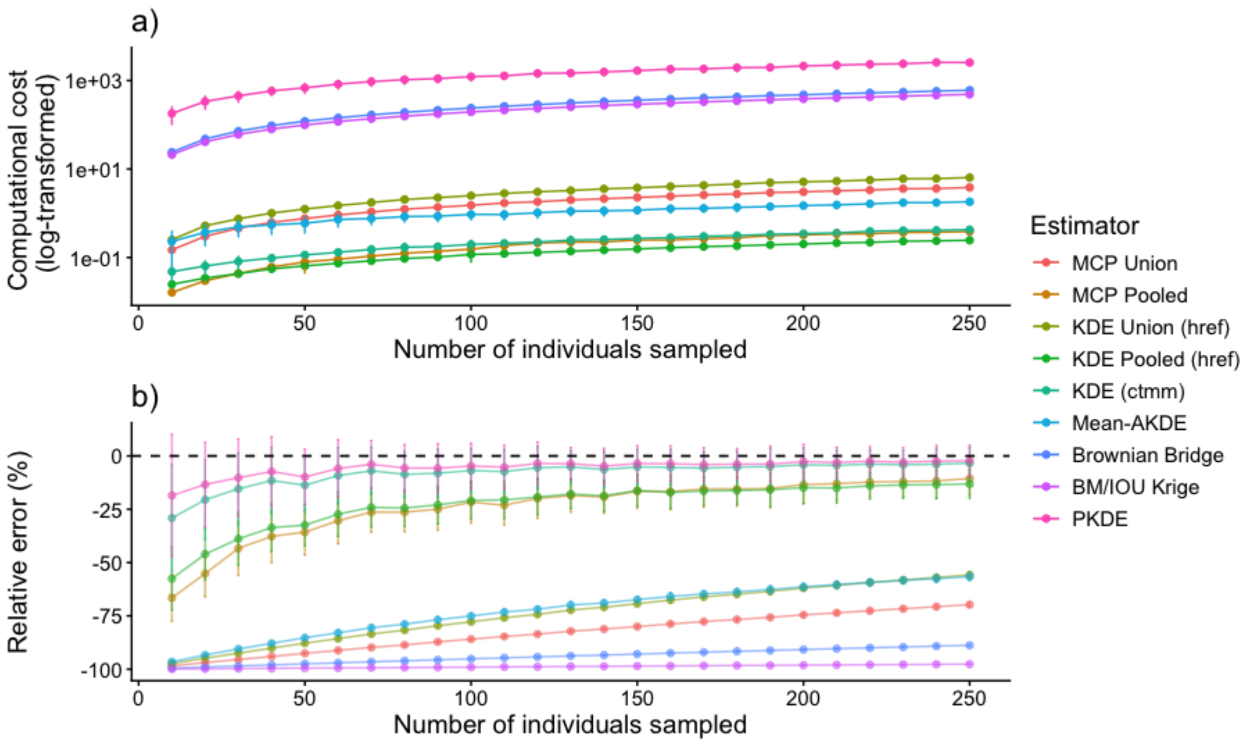


**Figure S7:** a) Computational cost in units of seconds (log-transformed) and b) mean relative error (%) of the population range area estimate for each method, as a function of the number of individuals sampled. Based on data simulated as specified in the main text, with 100 simulation runs per sample size. Error bars represent standard deviation. Negative relative error values on the y-axis in b) represent underestimation of the population range when compared to the true population range.

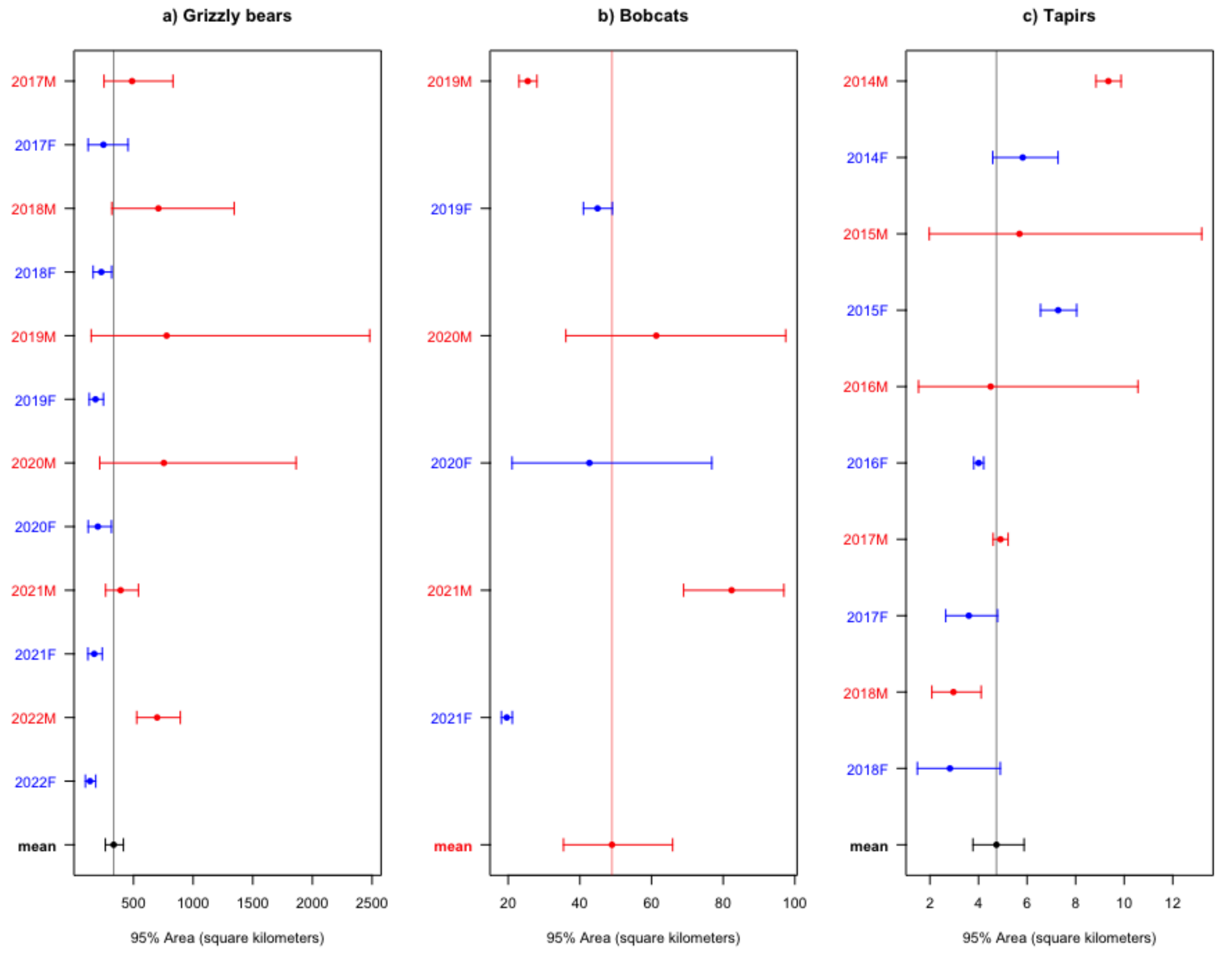


**Figure S8:** Yearly sex-specific differences in home range size for the three focal species: a) grizzly bears, b) bobcats, and c) tapirs, along with 95% confidence intervals. Male home range size is indicated in red and with the suffix 'M' after the given year on the y-axis, and female home range size in blue with the suffix 'F'. Significant differences between male and female home range size were observed for all three species.

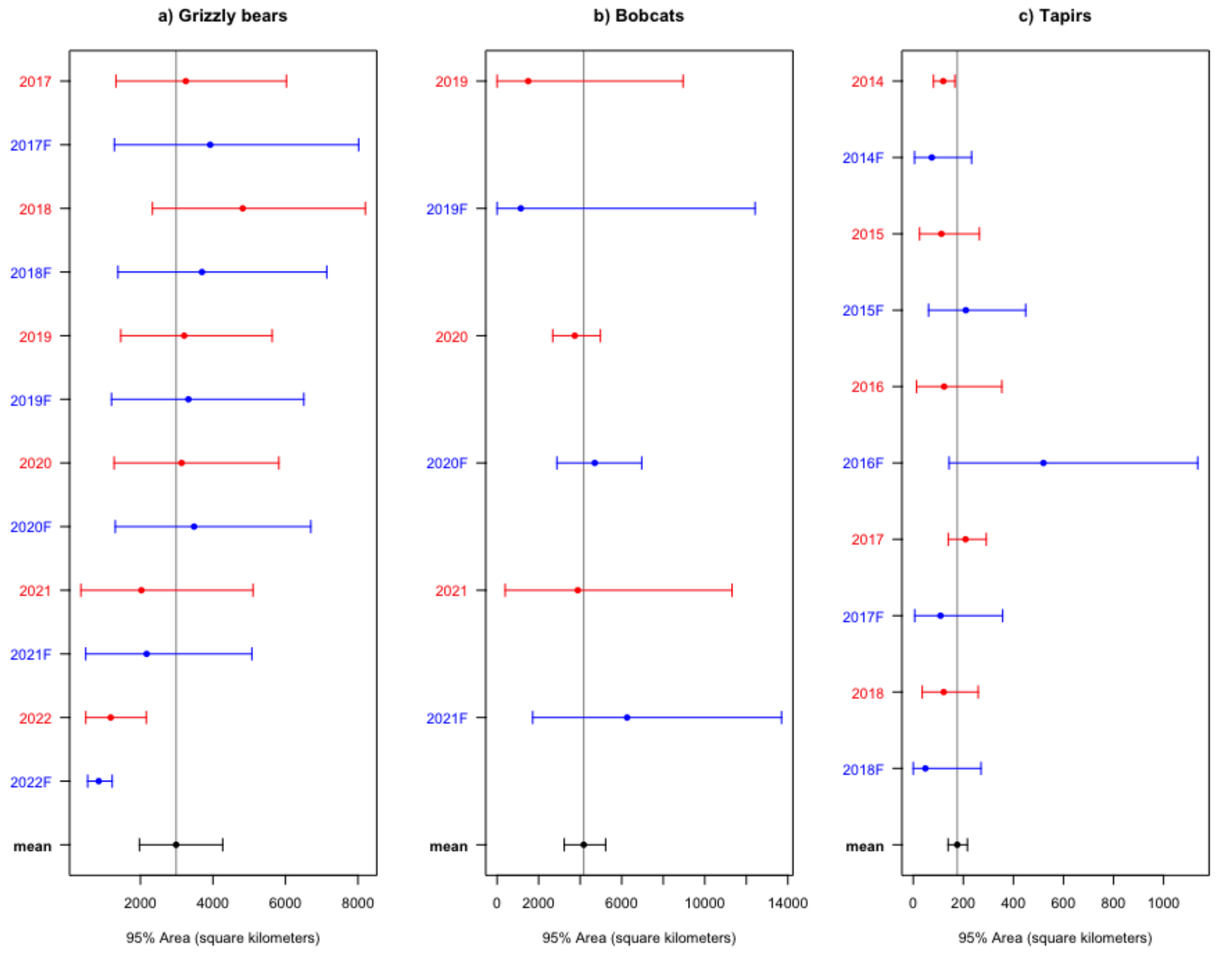


**Figure S9:** Yearly estimated PKDE population range size for the three focal species when considering all sampled individuals (indicated in red) vs. only females (indicated in blue, with the suffix 'F' following the given year on the y-axis). There is no significant difference in the population range size estimated by PKDE when all individuals are considered or just females.

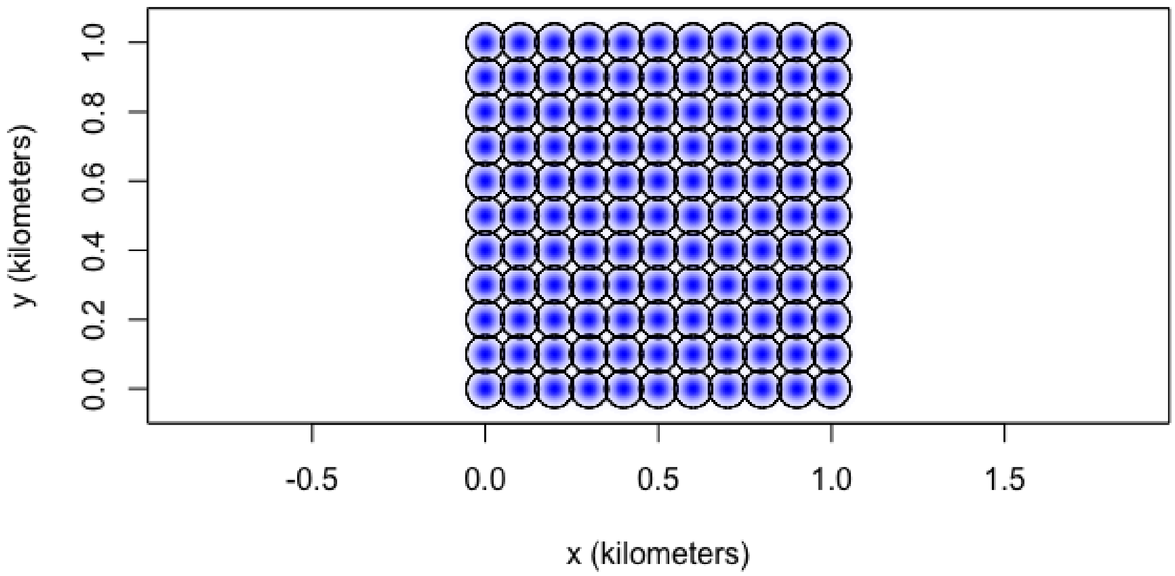


**Figure S10:** Simulation of a population where individuals have little to no overlap between their home ranges. The home range centers for the population consisting of 121 individuals are fixed in an 11x11 grid. Each individual has the same home range size, such that there is minimal overlap between individuals' home ranges. The black lines demarcate individual home ranges, and the blue shading corresponds to density estimates.

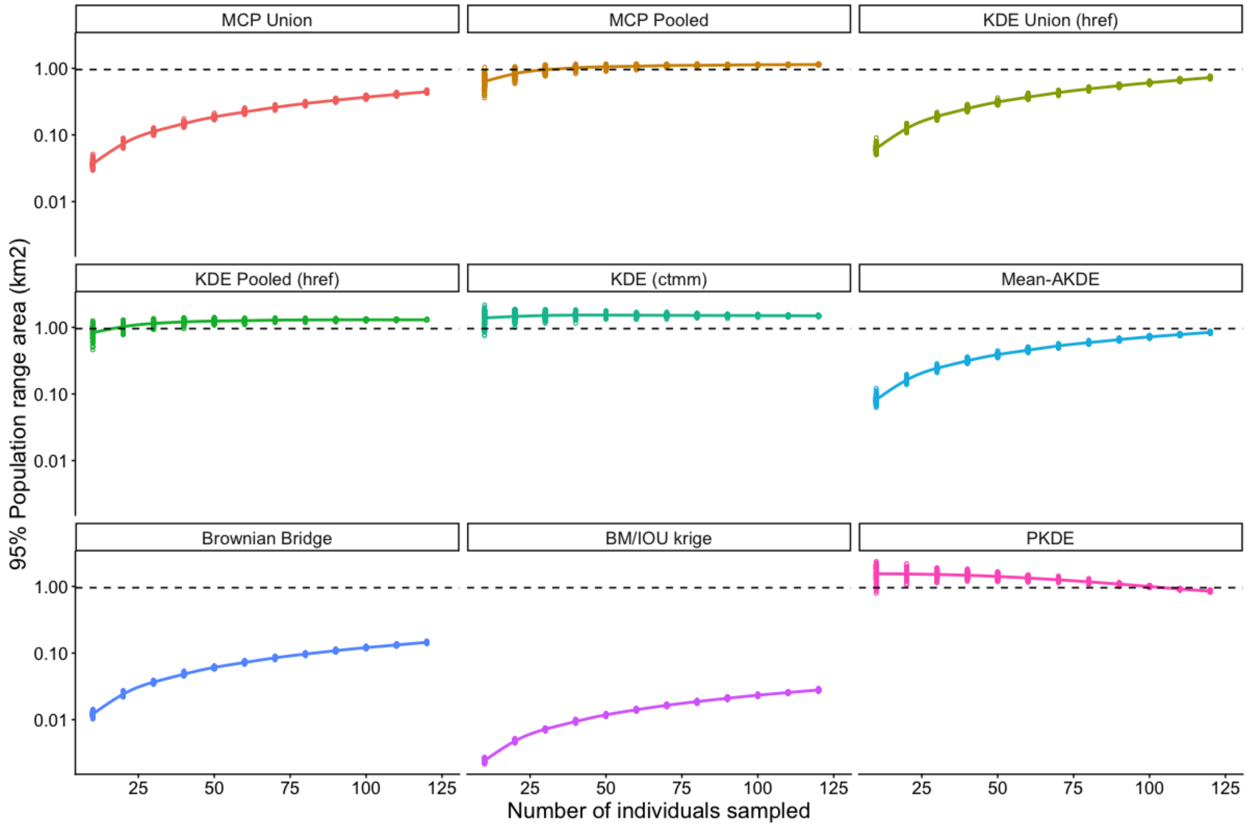


**Figure S11:** Validation of population space use estimators using simulated movement tracks where the individual home ranges have minimal overlap. Effect of sample size on the range (km$^2$) of a simulated population of 121 individuals predicted using MCP Union, MCP Pooled, KDE Union ($h_{ref}$), KDE Pooled ($h_{ref}$), KDE (*ctmm*), Mean-AKDE, Brownian Bridge, BM/IOU Krige, and PKDE. The y-axis is log-scaled. Dashed line indicates the true population range area. Each colored open circle represents one simulation run.
